# *Pseudomonas aeruginosa* inhibits *Rhizopus microsporus* germination through sequestration of free environmental iron

**DOI:** 10.1101/364877

**Authors:** Courtney Kousser, Callum Clark, Kerstin Voelz, Rebecca A. Hall

## Abstract

*Rhizopus spp* are the most common etiological agents of mucormycosis, causing over 90% mortality in disseminated infection. Key to pathogenesis is the ability of fungal spores to swell, germinate, and penetrate surrounding tissues. Antibiotic treatment in at-risk patients increases the probability of the patient developing mucormycosis, suggesting that bacteria have the potential to control the growth of the fungus. However, research into polymicrobial relationships involving *Rhizopus spp* has not been extensively explored. Here we show that co-culturing *Rhizopus microsporus* and *Pseudomonas aeruginosa* results in the inhibition of spore germination. This inhibition was mediated via the secretion of bacterial siderophores, confirming the essential role of iron for fungal growth. Addition of *P. aeruginosa* siderophores to *R. microsporus* spores in the zebrafish larval model of infection resulted in inhibition of fungal germination and reduced host mortality. Therefore, during infection antibacterial treatment may relieve bacterial imposed nutrient restriction resulting in secondary fungal infections.

## Introduction

Mucormycosis is a life threatening, disfiguring infection caused by ubiquitous environmental fungi belonging to the order *Mucorales*, with *Rhizopus spp*. accounting for approximately 70% of infections (Alvarez et al. 2009; Roden et al. 2005). In healthy individuals, innate immune cells are capable of controlling spore germination, thus preventing infection (Waldorf et al. 1984). However, patients with uncontrolled diabetes, cancer, neutropenia, burn/traumatic wounds, post-transplantation and those undergoing corticosteroid therapy or renal dialysis are prone to mucormycosis (Ibrahim et al. 2012; Spellberg et al. 2005). Mucormycetes are inherently resistant to antifungals, requiring surgical debridement of infected tissue followed by an aggressive antifungal regime. As a result, mucormycosis is associated with high mortality rates (up to 96% in disseminated infections), and significant morbidity (Roden et al. 2005).

Mucorales spores enter the body through inhalation or open wounds (Rammaert et al. 2012). As a result mucormycosis is commonly associated with pulmonary, rhinocerebral, or cutaneous infections (Torres-Narbona et al. 2007). Germination is key to the pathogenesis of Mucormycetes, leading to tissue penetration, endothelial angioinvasion, and vessel thrombosis, ultimately resulting in debilitating necrosis (Ibrahim et al. 2012). Traumatic and burn wound infections, including military-associated blast wounds, are known predisposing conditions for mucormycosis in the immunocompetent (Roden et al. 2005), with over 70% of these infections being polymicrobial in nature (Warkentien et al. 2015; Akers et al. 2014). *Pseudomonas aeruginosa*, *Staphylococcus aureus*, and *Escherichia coli* are the most commonly co-isolated bacterial species from chronic wounds (Gjødsbøl et al. 2006; Kalan et al. 2016), and are therefore likely to interact and compete with Mucorales spores. In addition, the emergence of mucormycosis has been associated with broad-spectrum antimicrobial treatment (Struck & Gille 2013; Baker 1957; Nash et al. 1971), suggesting that the surrounding microbiome plays a role in controlling fungal growth.

Here we show that *P. aeruginosa* inhibits the germination, and therefore virulence, of *Rhizopus microsporus*, a common cause of mucormycosis. This inhibition was mainly caused by bacterial secretion of iron-chelating molecules, which sequester iron from the fungus. Considering the prevalence of *P. aeruginosa* and *R. microsporus* in traumatic wounds, it is hypothesised that antimicrobial treatment of *P. aeruginosa* infections would reduce siderophore availability, increasing local iron concentrations promoting *R. microsporus* germination, and leading to fatal secondary Mucormycete infections.

## Results

### *Pseudomonas aeruginosa* strongly inhibits the germination of *Rhizopus microsporus*

Key to the pathogenesis of mucormycosis is the ability of spores to germinate and penetrate surrounding tissues (Ibrahim et al. 2012). To identify whether bacteria can influence fungal germination, *R. microsporus* spores were co-cultured with *Pseudomonas aeruginosa, Burkholderia cenocepacia*, *Staphylococcus aureus*, and *Escherichia coli*. Co-culture of *R. microsporus* with *P. aeruginosa* at multiplicities of infection (MOI) of 1:50 and 1:100 resulted in 56.8% (+/- 8.269, p = 0.0023) and 92% (+/- 2.784, p < 0.001) inhibition of fungal germination, respectively (Figure 1A, B). Conversely, co-culturing *R. microsporus* spores with *S. aureus, E. coli*, and *B. cenocepacia* did not affect fungal growth at any of the MOIs tested (Figure 1A). Taken together, the results obtained from live co-cultures between *R. microsporus* and *P. aeruginosa* indicates that these two microbes undergo a competitive relationship resulting in reduced fungal germination.

**Figure 1.**
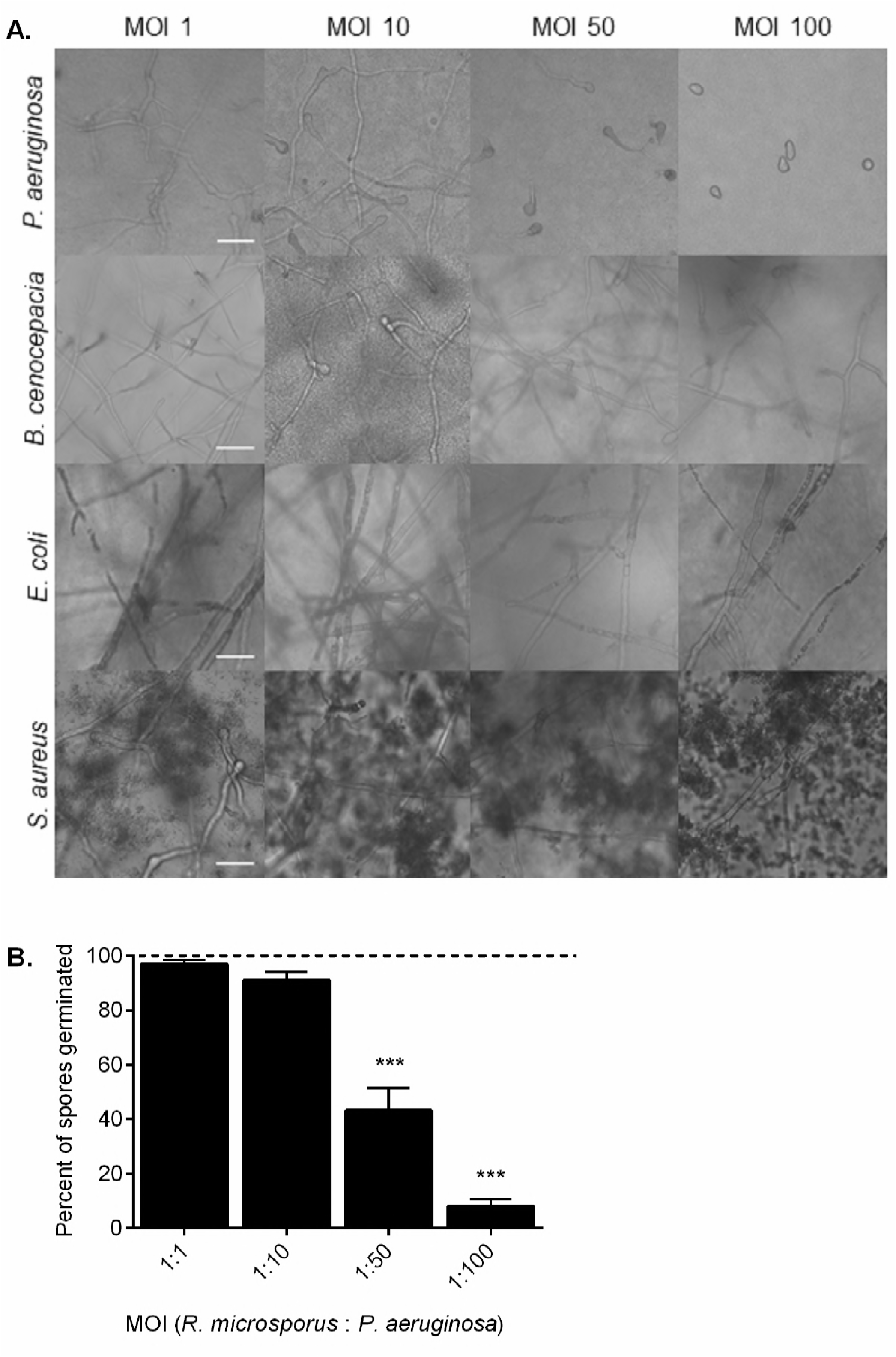
*Pseudomonas aeruginosa* strongly inhibits the germination of *Rhizopus microsporus*. *R. microsporus* spores were incubated with live *P. aeruginosa*, *B. cenocepacia, S. aureus*, and *E. coli* at increasing multiplicities of infection (MOI) for 24 h (A) Representative images after 24 h exposure (37°C, static). Scale bars depict 50 µm. (B) Per cent of spores germinated after 24 h exposure to *P. aeruginosa*. Dotted line represents spore-only control. One-way ANOVA performed on arcsine transformed data (n=6). *** = p < 0.001. Error bars depict SEM. Source files for all microscopy images used in the quantitative analysis are available in Figure 1-Source data 1.

### *P. aeruginosa* inhibits spore germination through secreted factors

Microbes are able to communicate through the secretion of secondary metabolites, quorum sensing molecules, and metabolic by-products (Jabra-Rizk et al. 2006; Peleg et al. 2008; Boon et al. 2008; Penner et al. 2016; Lopez-Medina et al. 2015; Hogan 2002). Therefore, to deduce whether the observed inhibition of *R. microsporus* germination was a result of direct cell-cell interactions or mediated through secreted products, *R. microsporus* spores were incubated in *P. aeruginosa* spent culture supernatants. Incubation of *R. microsporus* spores with 50% *P. aeruginosa* supernatant resulted in 94.4% (+/- 0.01769, p = 0.0022) inhibition of fungal growth (Figure 2A), confirming that the inhibitory molecule(s) are secreted by *P. aeruginosa*. Time-lapse microscopy confirmed that the presence of the supernatant resulted in a significant reduction in spore germination (Figure 2B, C, Video 1 and 2), with only 6.7% (+/- 3.8, p < 0.0001) of spores germinating after 18 hr. However, the inhibition of germination did not affect spore swelling (Video 1 and 2). To deduce whether *P. aeruginosa* supernatants are able to inhibit fungal growth after the initiation of germination, spores were pre-germinated, and then subsequently incubated with 50% supernatant. Fungal growth was significantly reduced (by 81.4% +/- 5.252, p = 0.0286) in the presence of the supernatant compared to the media control (Fig. 2D). Therefore, *P. aeruginosa* supernatants are able to inhibit both germination and growth of *R. microsporus*.

To determine whether the inhibitory molecule(s) is produced by other *P. aeruginosa* strains, we tested the ability of supernatants from a series of *P. aeruginosa* isolates to inhibit spore germination. *R. microsporus* germination was inhibited in the presence of supernatants from all *P. aeruginosa* clinical isolates (Fig. 2E), suggesting that the production of this inhibitory molecule is a general trait of *P. aeruginosa* and is not limited to laboratory-evolved strains.

**Figure 2.**
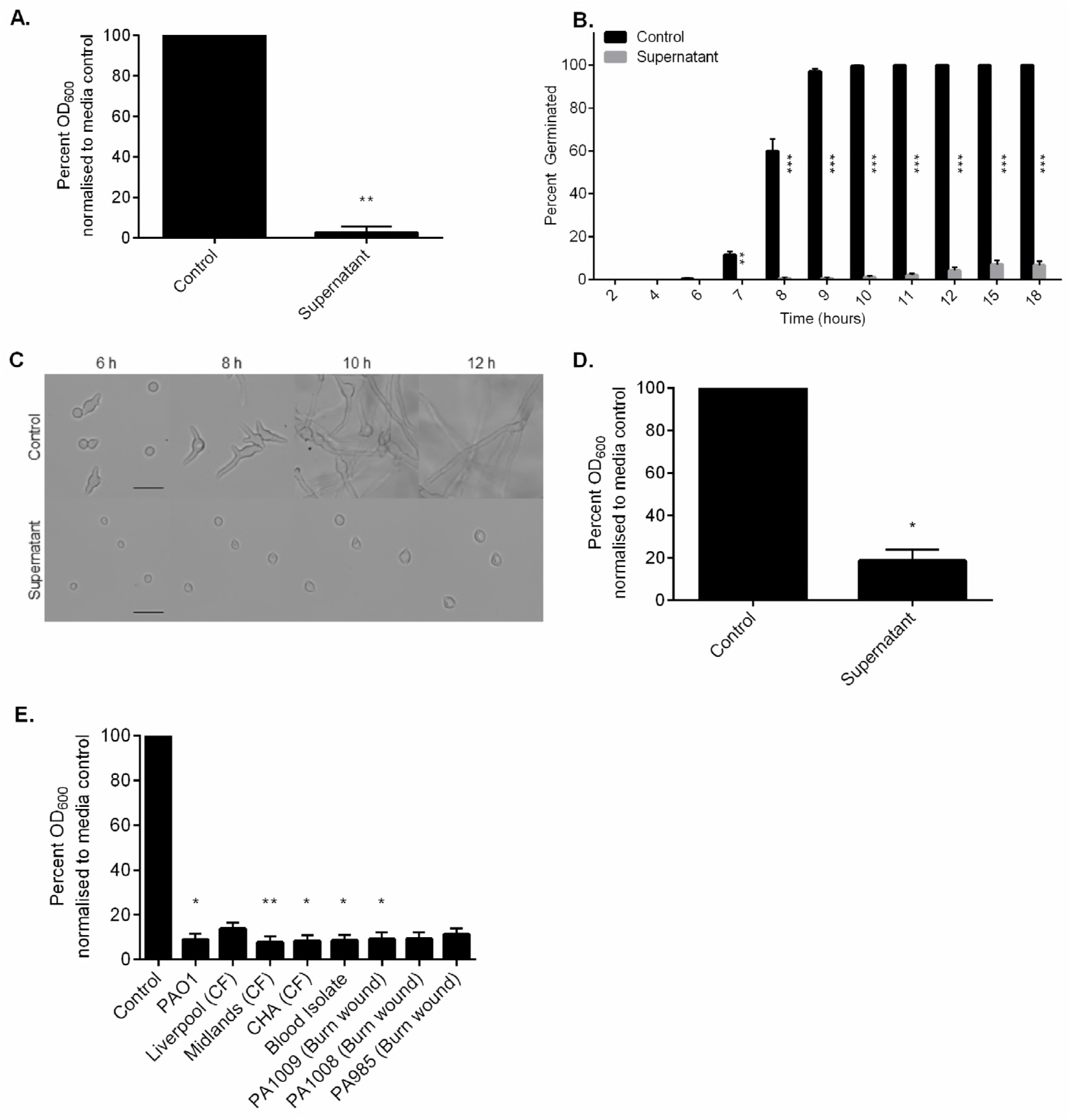
*P. aeruginosa* inhibits spore germination through secreted factors. *R. microsporus*spores were exposed to 50% *P. aeruginosa* PAO1 supernatant for 24 h. (A) Fungal growth was measured through absorbance (OD_600_) and normalised to media control (n=3). To determine the point of inhibition, spore germination was observed via live-cell imaging and (B) the per cent of spores germinated over time was quantified (n=4, Two-way ANOVA performed on arcsine transformed data). (C) Representative images of spores germinating over time were collected. Scale bar = 50 µm. (D) To determine whether the supernatant also inhibits the continuation of growth after germination is initiated, spores were incubated in SAB for 4-5 h until germlings emerged, and subsequently exposed to 50% PAO1 supernatant for 18 h (n=4, Mann-Whitney *U* test). (E) To test whether this is a lab strain-specific phenomenon, *R. microsporus* spores were exposed to supernatants from *P. aeruginosa* clinical isolates for 24 h (n=5). Fungal growth was determined through absorbance (OD_600_). All data was analysed by a Kruskal-Wallis test with Dunn’s multiple comparisons test unless indicated otherwise. * = p < 0.05, ** = p < 0.01, *** = p < 0.001. Error bars depict SEM. Source files for all microscopy movies used in the quantitative analysis are available in Figure 2 -source data 1.

As fungal germination is dependent on environmental pH and nutrient availability (Buffo et al. 1984; Singh et al. 2016), we assessed whether the addition of the supernatant was inhibiting germination through modulation of these parameters. Addition of the bacterial supernatant to SAB broth resulted in mild alkalisation of the media (pH 7.33 vs. 6.45). However, adjusting the pH of the control media to pH 7.33, to mimic the conditions in media containing the *P. aeruginosa* supernatant, did not affect *R. microsporus* germination rates (Figure 2 - figure supplement 1). To elucidate the role of macronutrient restriction, SAB broth was diluted with 50% phosphate buffered solution (PBS) to mimic the nutrient limitation imposed by the addition of 50% supernatant. However, the spores were still able to germinate under these conditions (Figure 2 - figure supplement 1). Therefore, *P. aeruginosa* secretes a molecule(s) that is able to inhibit *R. microsporus* germination independent of pH and nutrient limitation.

### Inhibition of *R. microsporus* germination is not mediated by quorum sensing molecules or pyocyanin

Bacteria secrete a diverse range of proteins and secondary metabolites to aid in host colonisation and inter-species competition. To determine whether the secreted factor responsible for inhibiting spore germination is proteinaceous, *P. aeruginosa* supernatants were boiled or treated with Proteinase K to degrade any secreted proteins. Supernatants that were boiled or treated with Proteinase K inhibited *R. microsporus* growth (97.62%, +/- 1.558, p = 0.0355 and 99.03%, +/- 1.634, p = 0.0140, respectively Figure 3A), suggesting that a secreted, heat-stable molecule mediates the observed inhibition of *R. microsporus* germination.

*P. aeruginosa* secretes several heat-stable cell density dependent signalling molecules into the environment to regulate virulence by sensing population density and inducing the expression or inhibition of population-dependent mechanisms (Waters & Bassler 2005). These quorum sensing molecules (QSMs) are well known to regulate intra- and inter-species interactions including inhibiting the morphological switch of *Candida albicans* (Enjalbert & Whiteway 2005; Davies 1998; Hogan et al. 2004; Cruz et al. 2013). Therefore, we tested the ability of the major *P. aeruginosa* QSMs to inhibit *R. microsporus* germination. Exposure of *R. microsporus* spores to N-butanoyl-l-homoserine lactone (C4 HSL), N-hexanoyl-DL-homoserine lactone (C6 HSL), and N-octanoyl-L-homoserine lactone (C8 HSL), did not affect fungal growth (Figure 3 – figure supplement 1). At high concentrations (200 μM) N-(3-oxododecanoyl)-L-homoserine lactone (C12 HSL) resulted in 42.1 % (+/- 0.1518, p =0.1331) reduction in fungal growth (Figure 3B). Therefore, secreted QSMs appear to not be the major regulators of *R. microsporus* growth.

Pyocyanin is a heat stable, secreted blue-pigmented toxin, which is known to increase the virulence of *P. aeruginosa* by depressing the host immune responses through induction of neutrophil apoptosis (Allen et al. 2005; Prince et al. 2013). Pyocyanin also inhibits the growth and morphogenesis of *C. albicans* and *Aspergillus fumigatus* (Lau et al. 2004). Therefore, we determined whether the presence of pyocyanin in the supernatant was inhibiting the germination of *R. microsporus*. Addition of purified pyocyanin resulted in 31.4% (+/- 0.1434, p > 0.9999) inhibition of *R. microsporus* growth at concentrations above 100 μM (Figure 3C). To deduce whether these pyocyanin concentrations were physiologically relevant the concentration of pyocyanin in the *P. aeruginosa* supernatants was quantified. Growth of *P. aeruginosa* in LB media at 200 rpm, 37°C did not result in the secretion of detectable levels of pyocyanin. Therefore, the inhibition of *R. microsporus* growth in the *P. aeruginosa* supernatant was not due to pyocyanin.

To determine whether the secreted factor is a lipophilic molecule, chloroform extractions were performed. The organic phase of the supernatant did not significantly inhibit spore germination (18% inhibition, +/- 3.058, p = 0.0926), while the aqueous phase maintained its inhibitory action (95.1% inhibition, +/- 0.9559, p = 0.0079, Figure 3D). Therefore, a secreted, heat-stable, water-soluble molecule(s) inhibits the growth of *R. microsporus*.

**Figure 3.**
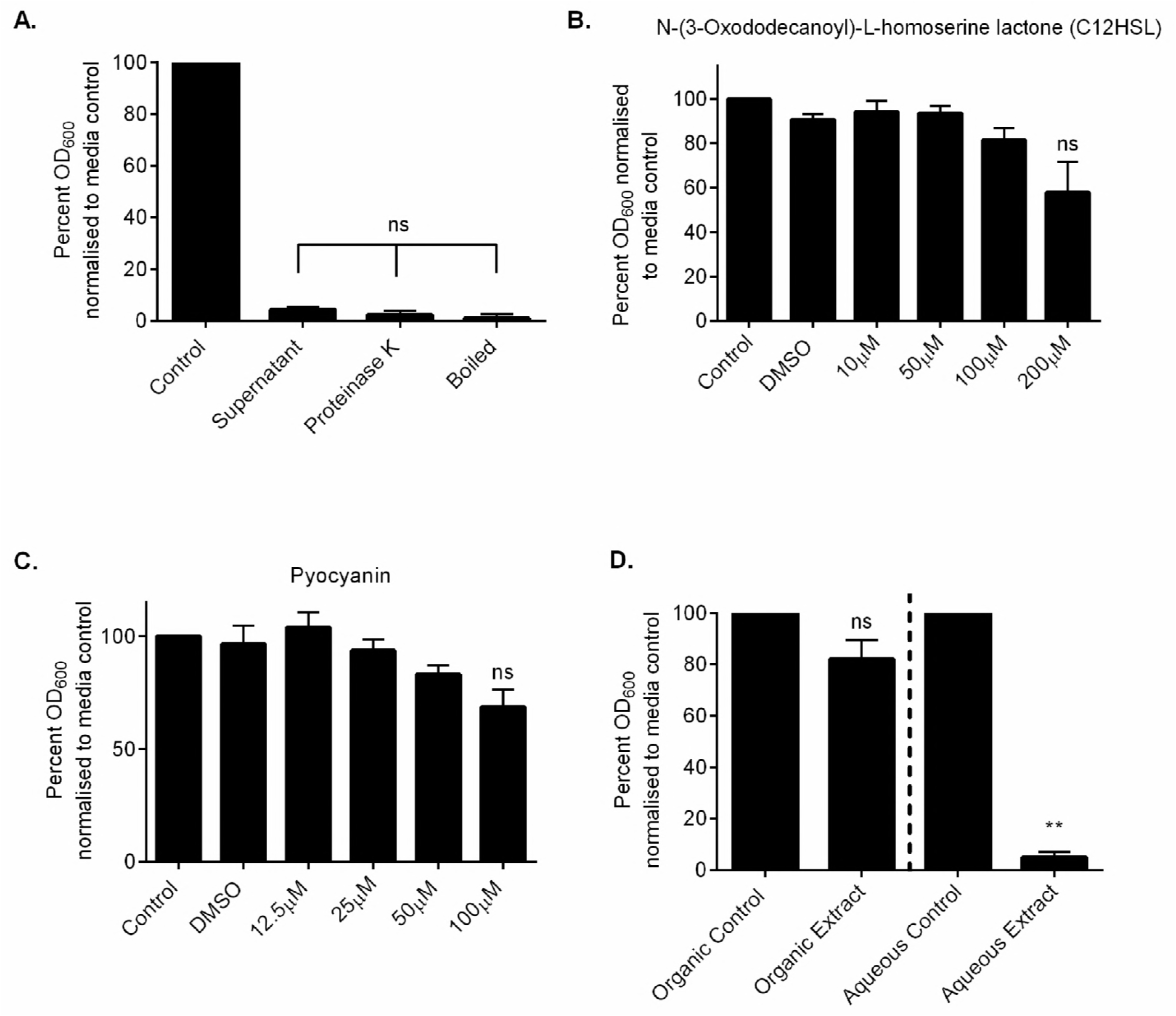
Inhibition of *R. microsporus* growth by *P. aeruginosa* secreted factors is mediated by a hydrophilic heat stable molecule. (A) PAO1 supernatant was treated with 150 µg/ml Proteinase K or boiled (100°C for 1 hour) to determine whether the inhibitory factor is a heat stable protein. Spores were exposed to treated supernatants and fungal growth was quantified through absorbance (OD_600_) after 24 h incubation at 37°C (n=6). (B) *R. microsporus* spores were incubated in SAB media for 24 h at 37°C with increasing concentrations of N-(3-oxododecanoyl)-Lhomoserine lactone (C12 HSL), and (C) pyocyanin. Fungal growth was quantified by absorbance (OD_600_) and normalised to media control (n=3). All data was analysed by a Kruskal-Wallis test with Dunn’s multiple comparisons test. Error bars depict SEM. (D) Chloroform extractions were performed to separate aqueous and organic molecules. *R. microsporus* spores were exposed to the aqueous and organic extracts for 24 h and fungal growth was determined through absorbance (OD_600_) (n=5). All data analysed with Kruskal-Wallis with Dunn’s multiple comparison’s test unless specified otherwise. Error bars depict SEM. ** = p < 0.01, *** = p < 0.001.

### *P. aeruginosa* inhibits *R. microsporus* germination via iron sequestration

Research has established the importance of metal micronutrients for microbial growth and pathogenicity, and the ability of host metal sequestering proteins to inhibit both fungal and bacterial growth through nutritional immunity (Weinberg 1975; Corbin et al. 2008; Foster 1939). Iron, zinc, copper, and manganese are considered the most important trace metals for the growth of fungi and the availability of iron is key to the pathogenesis of Mucormycetes (Ballou & Wilson 2016; Ibrahim et al. 2008). Therefore, we investigated whether the supernatants were imposing micronutrient restriction on *R. microsporus*. Supplementing the supernatants with iron was able to partially restore fungal growth, resulting in 46.2% *R. microsporus* growth at concentrations above 200 µM (+/- 6.660, Fig. 4A and C) and an insignificant difference as compared to the control (p > 0.9999). However, supplementation with zinc, copper or manganese was unable to rescue *R. microsporus* growth (Figure 4 - figure supplement 1). Therefore, *P. aeruginosa* supernatants specifically sequester iron from the environment, resulting in the inhibition of *R. microsporus* growth. To confirm the inhibition of spore germination observed in the co-cultures was also attributed to iron restriction, co-cultures of *R. microsporus* and *P. aeruginosa* were spiked with iron. Germination and therefore growth of *R. microsporus* in the co-culture was recovered at concentrations above 100 µM (41.9% +/- 12.46, p = 0.1540) to levels comparable to those observed for iron spiked supernatants (Figure 4B), confirming that in both scenarios *P. aeruginosa* sequesters iron, inhibiting the growth and germination of *R. microsporus*.

Iron starvation has previously been shown to up-regulate the high affinity iron permease *FTR1* in other *Rhizopus* species (34). Therefore, to confirm that *R. microsporus* is undergoing iron starvation in the presence of *P. aeruginosa* supernatants, the expression levels of *FTR1* were determined by qRT-PCR. *FTR1* was highly upregulated (100-fold increase) when exposed to 50% *P. aeruginosa* supernatant for 7 hr, as compared to the control (Figure 4D). This confirms that *P. aeruginosa* mediated iron restriction inhibits *R. microsporus* growth and germination.

Iron stress has been shown to induce apoptosis in *R. oryzae* after prolonged starvation (Shirazi et al. 2015). Therefore, if spores are undergoing iron starvation when exposed to *P. aeruginosa* supernatant, prolonged exposure should decrease survival. To isolate the effects of the supernatant, we used 100% *P. aeruginosa* supernatant to monitor spore survival over time. In this condition, the viability of spores was reduced by 82.40% (+/- 13.44) after 24 hr, and no viable spores were recovered after 120 hr (p = 0.0490, Figure 4E). This indicates that iron is essential for the survival and pathogenicity of *R. microsporus*.

**Figure 4.**
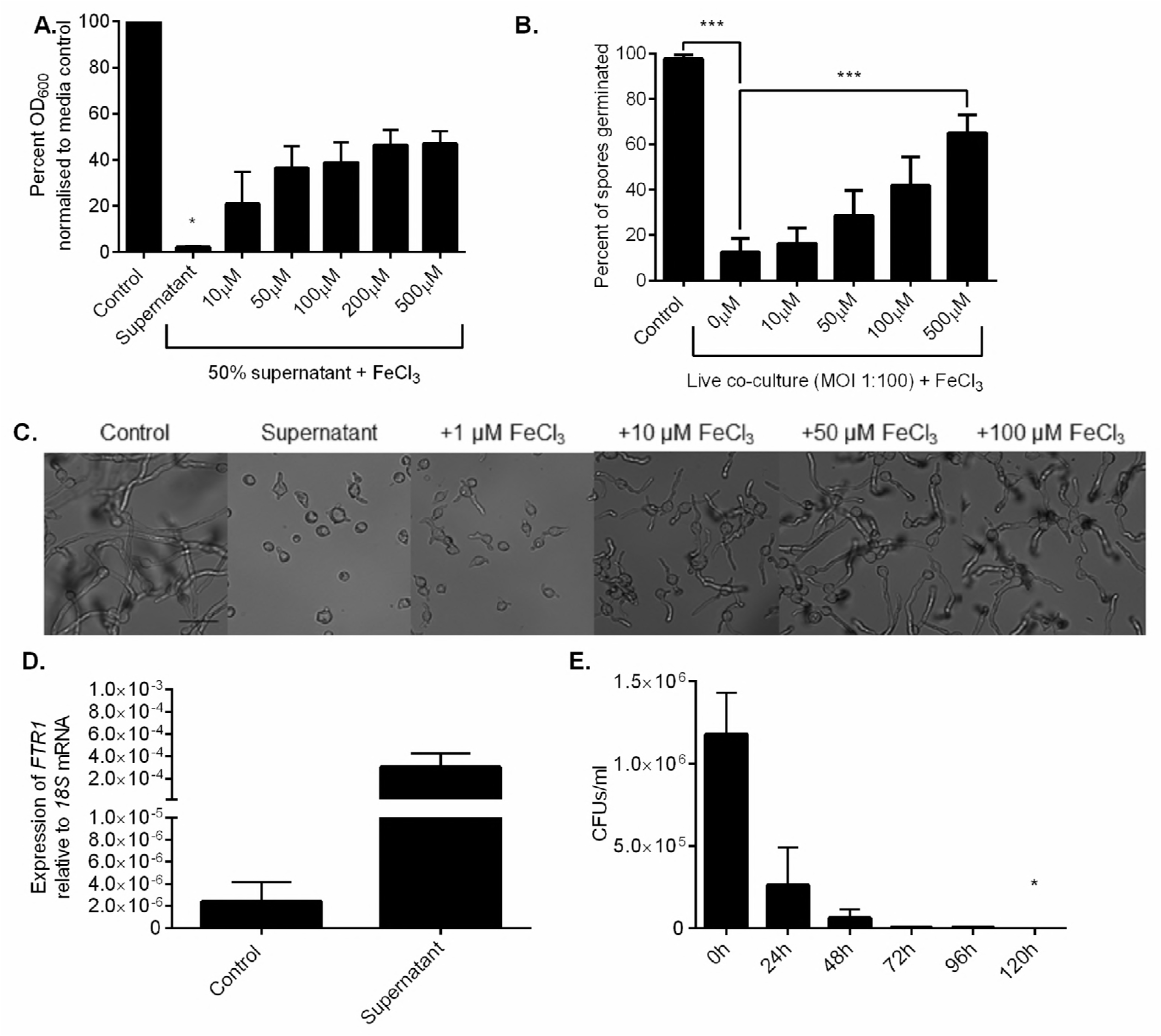
*P. aeruginosa* inhibits*R. microsporus* germination via iron sequestration. *R. microsporus* spores were exposed to (A) 50% *P. aeruginosa* supernatant and spiked with increasing concentrations of iron chloride for 24 h statically at 37°C. Fungal growth was measured through absorbance (OD_600_) and normalised to media control (n=3, Kruskal-Wallis test with Dunn’s multiple comparisons test). (B) This ability to rescue was confirmed in a live co-culture setting, where the addition of exogenous iron increased the per cent of spores germinated after 24 h in a dose-dependent manner (n=8). As the addition of iron in 50% supernatant increased overall growth, (C) representative images were collected at 9 h to confirm ability to rescue germination. Scale bar depicts 50 µm. (D) Iron starvation of *R. microsporus* spores after 7 h exposure to *P. aeruginosa* supernatant was determined through strong upregulation of the high-affinity iron permease *FTR1* (n=2). (E) As iron starvation is associated with mucormycete apoptosis, the viability of spores exposed to 100% *P. aeruginosa* supernatant over time was quantified by counting colony forming units (CFUs) every 24 h for 120 h (n=3, Kruskal-Wallis test with Dunn’s multiple comparisons test). * = p < 0.05, *** = p < 0.001. Source files for all microscopy images used in the quantitative analysis are available in Figure 4-source data 1.

To delineate whether bacteria-associated iron restriction inhibits the growth of Mucormycetes in general, we tested the ability of *P. aeruginosa* supernatant to inhibit the growth of *R. microsporus var. microsporus, R. microsporus var. chinensis, R. delemar, and Mucor circinelloides*. The growth of all isolates was significantly reduced in the presence of *P. aeruginosa* supernatant [83.85% (+/- 15.23), 98.26% (+/- 1.977), 99.01% (+/- 0.8695), and 87.54% (+/- 10.78), respectively], and was significantly rescued by the addition of 100 μM of iron [64.36% (+/- 4.450), 41.30% (+/- 10.48), 55.14% (+/- 10.01), and 87.54% (+/- 6.219), respectively], confirming that this bacteria-associated inhibition of growth is a general trait of Mucormycetes (Figure 5).

**Figure 5.**
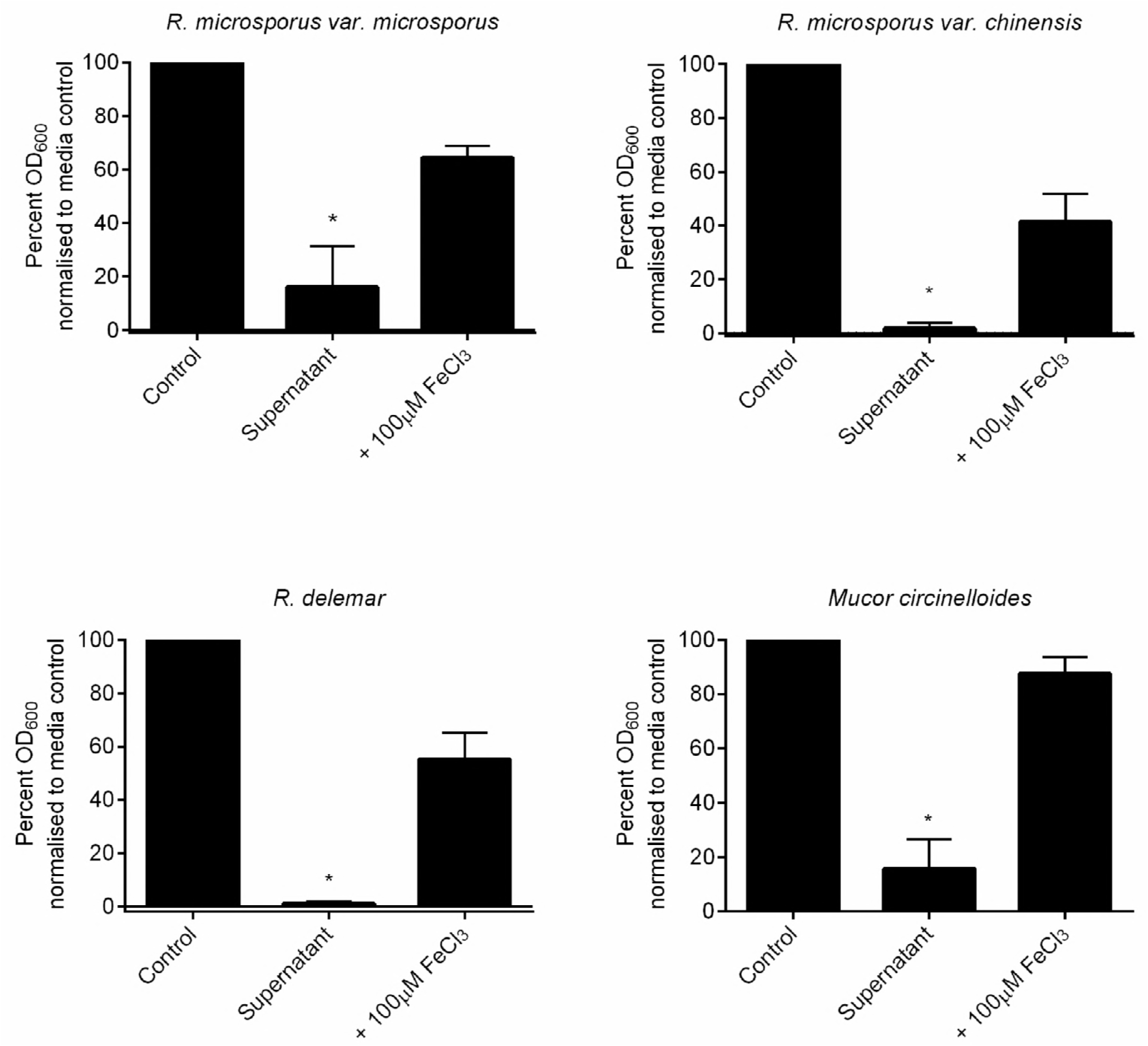
Iron-dependent inhibition of Mucormycetes by *P. aeruginosa* is not *R. microsporus* strain-specific. Most experiments in this study were performed using an *R. microsporus* clinical isolate. To ensure the inhibitory effect of *P. aeruginosa* is not limited to this isolate, *R. microsporus var. microsporus, R. microsporus var. chinensis, R. delemar*, and *Mucor circinelloides* were exposed to 50% *P. aeruginosa* supernatant with and without the addition of 100 µM FeCl_3_. Fungal growth was determined at 24 h by measuring absorbance (OD_600_) and normalising to control (n=3). * = p < 0.05. All data was analysed by a Kruskal-Wallis test with Dunn’s multiple comparisons test.

### Siderophore deficient *P. aeruginosa* rescues the growth of *R. microsporus*

Iron sequestering is mediated via iron binding proteins and molecules known as siderophores. Pyoverdine and pyochelin are the two predominate siderophores produced by *P. aeruginosa*, with pyoverdine exhibiting the highest affinity for iron (Braud, Hannauer, et al. 2009; Braud, Hoegy, et al. 2009). As the aqueous fraction of the bacterial supernatant was sufficient to inhibit fungal growth, we quantified fungal growth in the presence of the aqueous fractions of supernatants from *P. aeruginosa* strains deficient in both siderophores. Supernatants from siderophore defective *P. aeruginosa* strains (ΔpchEFΔpvdD) did not significantly inhibit fungal growth (Figure 6A), confirming that the iron restriction is mediated by siderophores.

To investigate whether pyoverdine alone is sufficient to inhibit *R. microsporus* growth, spores were exposed to exogenous pyoverdine (20-160 µg/ml). Incubation of fungal spores with pyoverdine significantly reduced fungal growth (79.4%, +/- 5.503, p = 0.0384, Figure 6B). To determine whether these concentrations of siderophores are physiologically relevant, the concentration of siderophores in the *P. aeruginosa* culture supernatants was quantified. Growth of *P. aeruginosa* in LB media resulted in the secretion of 58.9 µg/ml (+/- 1.194) siderophores. Therefore, the siderophore concentration in the supernatant is sufficient to inhibit *R. microsporus* growth and germination.

### Production of pyoverdine by *P. aeruginosa* is enhanced in the presence of *R. microsporus*

*C. albicans* can decrease *P. aeruginosa* siderophore production through suppression of the pyoverdine and pyochelin biosynthetic pathways (Lopez-Medina et al. 2015). To determine whether *R. microsporus* is also able to interfere with *P. aeruginosa* siderophore production, the concentration of pyoverdine during mono and co-culture was quantified. Surprisingly, the concentration of pyoverdine in co-cultures was 1.7 fold (+/- 0.47) higher in the co-cultures (MOI 1:100) compared to *P. aeruginosa* monoculture (Figure 6C). Therefore, the presence of *R. microsporus* enhances the production of pyoverdine, an essential virulence factor in *P. aeruginosa* infections.

### The concentration of siderophores produced by bacteria correlates with inhibition of *R. microsporus* growth

While live *E. coli, B. cenocepacia*, and *S. aureus* did not inhibit the growth of *R. microsporus*, these bacteria all produce siderophores (Courcol et al. 1991; O’Brien, I et al. 1970; Darling et al. 1998). To determine the ability of secreted factors to inhibit growth, *R. microsporus* spores were exposed to sterile supernatants from *E. coli, B. cenocepacia, Klebsiella pneumoniae* and *S. aureus* to determine their ability to inhibit *R. microsporus* growth as compared to *P. aeruginosa* PAO1 and PA14. Consistent with the co-culture experiments, *P. aeruginosa* was the only supernatant able to significantly inhibit growth (Figure 6D). We further investigated whether this lack of inhibition was associated with insufficient production of iron binding molecules by measuring the amount of siderophores produced after 24 hr growth in LB. There was a negative correlation between fungal growth and siderophore production (p = 0.0029), with the exception of *E. coli* (Figure 6E). This was surprising, as *E. coli* produces enterobactin, a siderophore with a high affinity (10^52^ M) for iron (Carrano & Raymond 1979). In agreement with this data, exogenous enterobactin did not inhibit *R. microsporus* growth (91%, +/- 6.170, p = 0.936, Figure 6F). Therefore, our data suggests that *R. microsporus* may utilise enterobactin as a xenosiderophore.

**Figure 6.**
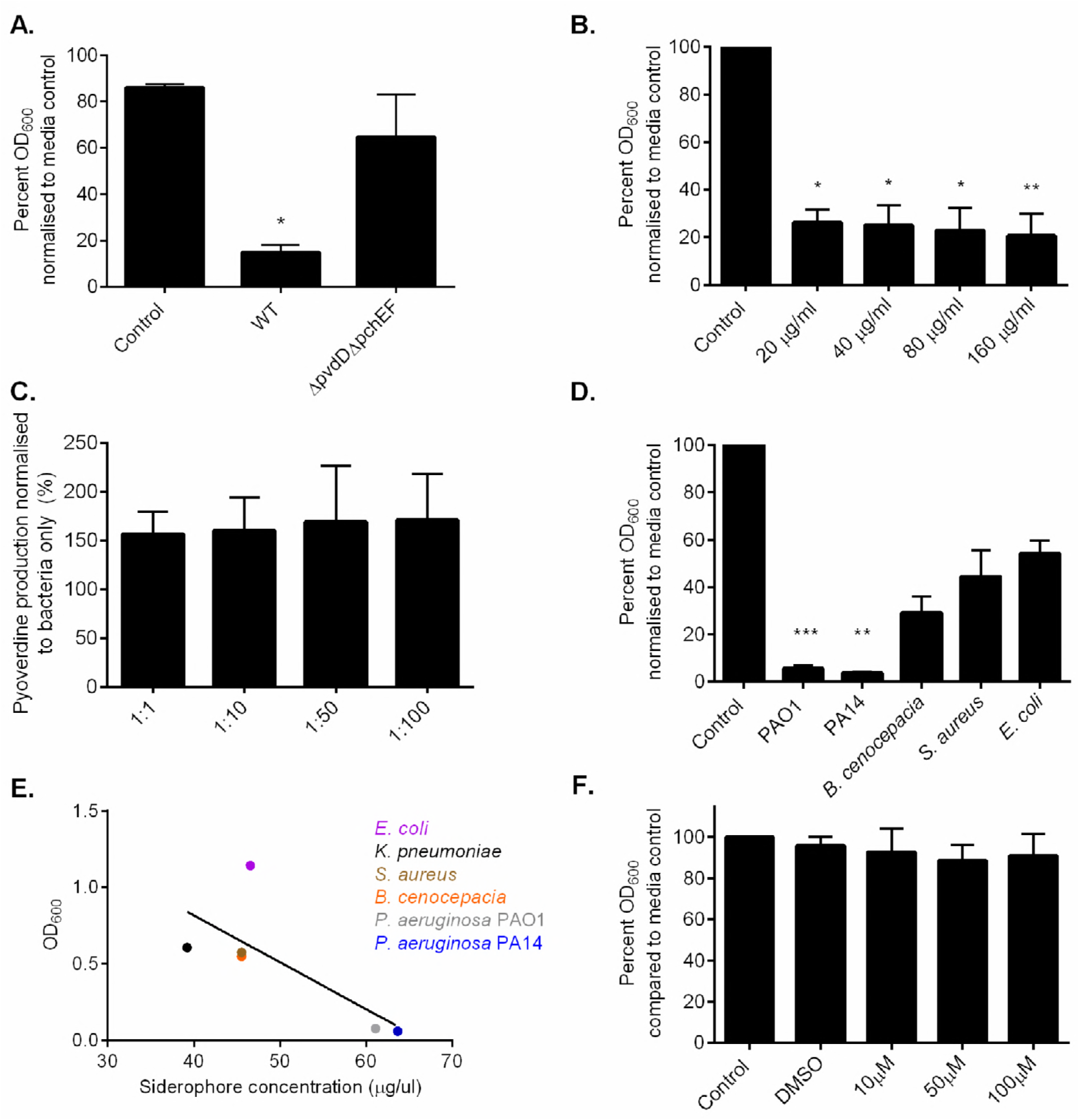
The ability of bacteria to inhibit growth of *R. microsporus* is associated with siderophore production. (A) Amberlite^®^ XAD-2^®^ was used to separate the aqueous portion of LB broth, WT, and siderophore mutant *P. aeruginosa* PAO1 (ΔpchEFΔpvdD) to isolate the fragment containing pyoverdine. *R. microsporus* spores were exposed to 50% of these extracts for 24 hr and fungal growth was determined via absorbance (OD_600_) and normalised to growth in SAB/whole LB (n=3). (B) *R. microsporus* spores were incubated in SAB with increasing concentrations of exogenous pyoverdine at 37°C for 24 h. Fungal growth was measured through absorbance (OD_600_) and normalised to media control (n=6) (C) *P. aeruginosa* was exposed to *R. microsporus* at a range of MOIs for 24 h (37°C). Pyoverdine production was measured through absorbance (405 nm) of co-culture supernatants and compared to 24 h *P. aeruginosa* mono-culture at the same concentration (n=4). (D) *R. microsporus* spores were exposed to 50% supernatants harvested from *P. aeruginosa* PAO1, *P. aeruginosa* PA14, *B. cenocepacia, S. aureus*, and *E. coli* for 24 h. Fungal growth was determined by absorbance (OD_600_) and normalised to control. (n=6). (E) Concentration of siderophores produced by bacteria was determined by using the Siderotec Assay (EmerginBio). Correlation between total siderophore production and fungal growth was determined by performing a linear regression with Pearson correlation (n=3). (F) *R. microsporus* spores were exposed to varying concentrations of purified enterobactin for 24 h at 370C (n=3). Fungal growth was determined by absorbance (OD_600_) and normalised to control. All data was analysed by a Kruskal-Wallis test with Dunn’s multiple comparisons test unless indicated otherwise. * = p < 0.05, **= p < 0.01, *** = p < 0.001.

### The bacterial siderophore, pyoverdine, inhibits *R. microsporus* virulence

To determine whether the effects of the bacterial siderophores have a role in controlling fungal infection in the host, we utilised the zebrafish larval model (Figure 7A). Co-injection of *R. microsporus* with pyoverdine (80 µg/ml) resulted in a mild but significant increase in fish survival when compared to spores alone (Figure 7B) with 89% (+/- 6.377) of fish surviving across a 96 hr time course. Together these data confirm that the presence of the *Pseudomonas spp*. high affinity siderophore, pyoverdine, is sufficient to inhibit the pathogenesis of *R. microsporus* and significantly decrease host damage.

**Figure 7.**
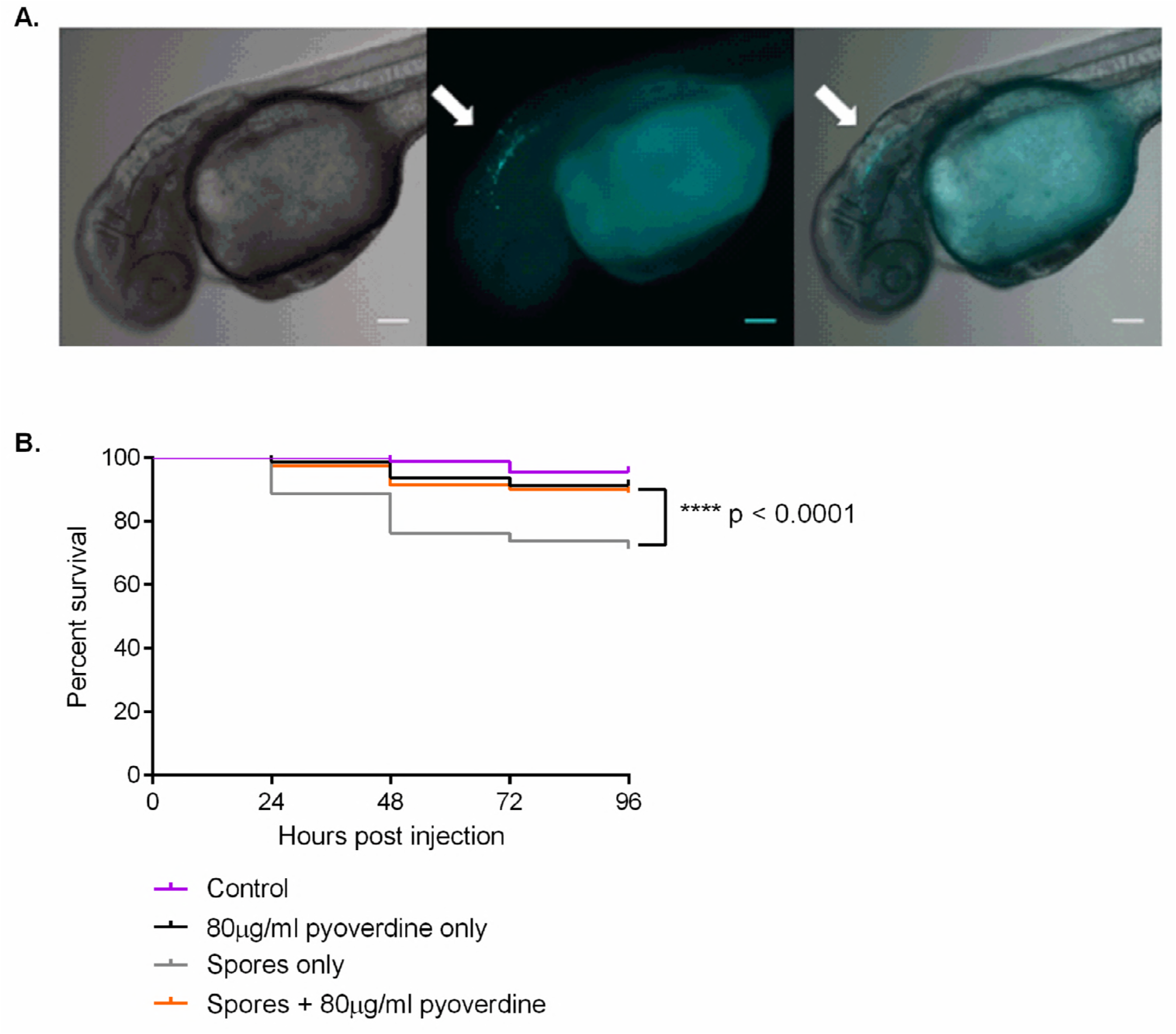
The bacterial siderophore, pyoverdine, inhibits *R. microsporus* virulence. To determine the impact of pyoverdine on fungal virulence within a host, zebrafish larvae were injected in the hindbrain with 50 spores +/- 80 μg/ml pyoverdine. (A) Representative images of zebrafish larvae at 0 hpi. White arrows indicate *R. microsporus* spores (Calcofluor White stain, cyan pseudo-coloured) located within hindbrain compartment. Scale bars depict 100 μm. (B) Survival of larvae was quantified over time. Shown are data pooled from four separate experiments with a total of 87, 80, 80, and 81 fish for control, pyoverdine only, spores only, and spores + pyoverdine, respectively. Data analysed with Mantel-Cox log-rank test.

## Discussion

Mucormycosis is a lethal infection with high mortality rates and lack of treatment options due to intrinsic antifungal resistance (Sun et al. 2002). Our current understanding of the pathogenesis is incomplete, especially when compared to other opportunistic fungal pathogens such as *C. albicans* and *A. fumigatus*. Because of this, it is important to understand the pressures Mucormycetes encounter within the human body. This not only includes pressures from the host, but also from the microbiota. Here we identify that *P. aeruginosa* is able to inhibit the germination of *R. microsporus* through the secretion of siderophores.

*Pseudomonas* species interact with and control the growth of a variety of fungal species including important plant and animal pathogens (Mowat et al. 2010; Hogan et al. 2004; Wallace et al. 2018). These interactions have been linked to a variety of contact dependent (Hogan & Kolter 2002) and bacterial secreted factors (Hogan et al. 2004; Mowat et al. 2010; Wallace et al. 2018). The most characterised secreted molecules known to affect fungal growth and morphology are the phenazines (Morales et al. 2013; Briard et al. 2015) and the homoserine lactones (Hogan et al. 2004; Mowat et al. 2010). For example, in *C. albicans* low levels of phenazines inhibit filamentation and biofilm formation, and are fungicidal at high concentrations (Gibson et al. 2009). Furthermore, the quorum sensing molecule, 3-oxo-C12-homoserine lactone (C12 HSL) induces apoptosis in *C. albicans* (Hogan et al. 2004) and *A. fumigatus* (Mowat et al. 2010). Despite this, these molecules had negligible impact on the growth of *R. microsporus*, confirming that other bacterial secreted factors control the growth of *R. microsporus*. However, high concentrations of C12 HSL (200 µM) had a marginal effect on *R. microsporus* growth, indicating that intra-species QS may play a role in polymicrobial biofilms.

Instead we identified that this antagonistic relationship between *P. aeruginosa* and *R. microsporus* to be the result of competition for iron. Iron acquisition is key to Mucorales pathogenesis (Ibrahim et al. 2010). For example, medical conditions (i.e. diabetic ketoacidosis) that result in increased serum levels of iron predispose individuals to Mucormycete infection (Artis et al. 1982), whereas iron chelation therapy, or reduction in fungal iron acquisition mechanisms reduce mortality in murine models of mucormycosis (Ibrahim et al. 2007; Ibrahim et al. 2010). *P. aeruginosa* secretes several iron binding molecules, with pyoverdine being the major siderophore with a high affinity for iron. In agreement with this, we found that exogenous pyoverdine, at concentrations equal to those secreted by *P. aeruginosa* in our culture conditions, was sufficient to inhibit the growth of *R. microsporus* to levels comparable to the bacterial supernatant. In addition, deletion of key enzymes in the biosynthesis pathways of the major *P. aeruginosa* siderophores was sufficient to reduce the effect of the bacterial supernatant, confirming a role for these siderophores in controlling fungal growth.

The presence of fungi has been shown to modulate the expression of siderophore biosynthetic genes in *P. aeruginosa*. For instance, *Candida albicans* was described as down-regulating the production of pyoverdine and pyochelin through secreted proteins (Lopez-Medina et al. 2015). Conversely, this study has found the production of pyoverdine to be increased in response to *P. aeruginosa* co-cultured with *R. microsporus*. This is clinically important, as pyoverdine production is directly linked to virulence of *P. aeruginosa* and is shown to modulate the production of other toxins (Meyer et al. 1996; Lamont et al. 2002).

The addition of exogenous iron or the inhibition of bacterial siderophore production only resulted in the restoration of approximately 50% fungal growth compared to media only controls, suggesting that other factors also contribute to this inhibition. However, in *Rhizopus oryzae*, iron starvation induces apoptosis (Shirazi et al. 2015), suggesting that spore viability may also be affected. In agreement with this, grown of *R. microsporus* in 100% *P. aeruginosa* supernatant decreased spore viability. As such, it is possible that reduced viability may account for the inability to completely rescue fungal growth. Interestingly though, exogenous iron was able to fully restore the growth of *Mucor cicinelloides*, suggesting that *R. microsporus* may undergo another species-specific interaction with *P. aeruginosa*.

*R. microsporus* can utilise some bacterial siderophores as sources of iron within the host, such as deferoxamine (a siderophore produced by some actinomycetes) to promote its growth and virulence. (Boelaert et al. 1993). However, unlike deferoxamine, and potentially entrobactin, *R. microsporus* cannot scavenge iron from pyoverdine, which suggests that molecules with similar structure may have the potential to be used to control mucormycosis. While utilising pyoverdine itself would be problematic due to its ability to enhance *P. aeruginosa* virulence (Lamont et al. 2002), this siderophore could provide a starting point for the development of novel iron chelators. Given that pyoverdine has also been shown to limit the growth of other invasive fungi, such as *A. fumigatus* (Penner et al. 2016; Sass et al. 2018), molecules based on pyoverdine may have wide implications for the treatment of a range of invasive fungal diseases. This is further enforced by the fact that the presence of pyoverdine in our zebrafish larval model of infection was able to reduce mortality. Similar effects have also be observed in mouse models of infection where deferasirox protects against mycormycosis (Ibrahim et al. 2007). Therefore, iron chelation therapy could be an important preventative treatment for mucormycosis.

Taken together, our results agree with the current understanding of Mucormycete pathogenesis where iron availability is considered essential for pathogenesis. However, here we present this in a different scenario where iron availability is controlled by surrounding bacteria. Given that a high percentage of invasive mucormycosis results from burn and blast wound infections, where iron availability will be high due to tissue damage, we propose that opportunistic bacteria like *P. aeruginosa* will sequester iron away from the fungus restricting fungal growth. In agreement with this, burn wound exudate enhances *P. aeruginosa* siderophore production (Gonzalez et al. 2016) resulting in high concentrations of pyoverdine in the wound. However, antibiotic treatment would reduce this competition for iron, and promote fungal germination. This, coupled with natural immunosuppression following trauma could lead to aggressive secondary mucormycosis (Kimura et al. 2010). Therefore, patients that have potentially been exposed to fungal spores (i.e. soldiers with blast wounds where significant environmental contamination of the wound has occurred) should be closely monitored for secondary fungal infections. The discovery of suitable iron chelators that do not promote bacterial virulence would be advantageous in this setting to help prevent fungal infection.

## Material and Methods

### Ethics

Zebrafish care and experiments were performed under Home Office project license 40/3681 and personal license I5B923969 in accordance with the Animal Scientific Procedures Act 1986.

### Strains and culture conditions

All media and chemicals were purchased from Sigma-Aldrich unless stated otherwise. For details of fungal and bacterial strains used, please see (Table 1). *R. microsporus* was routinely sub-cultured and maintained on Sabouraud 4% dextrose agar (SAB, Merck Millipore, Germany) and incubated for 10-14 days before use (25°C). Bacteria were maintained on Lysogeny broth (LB) with 2% agar.

**Table 1.**
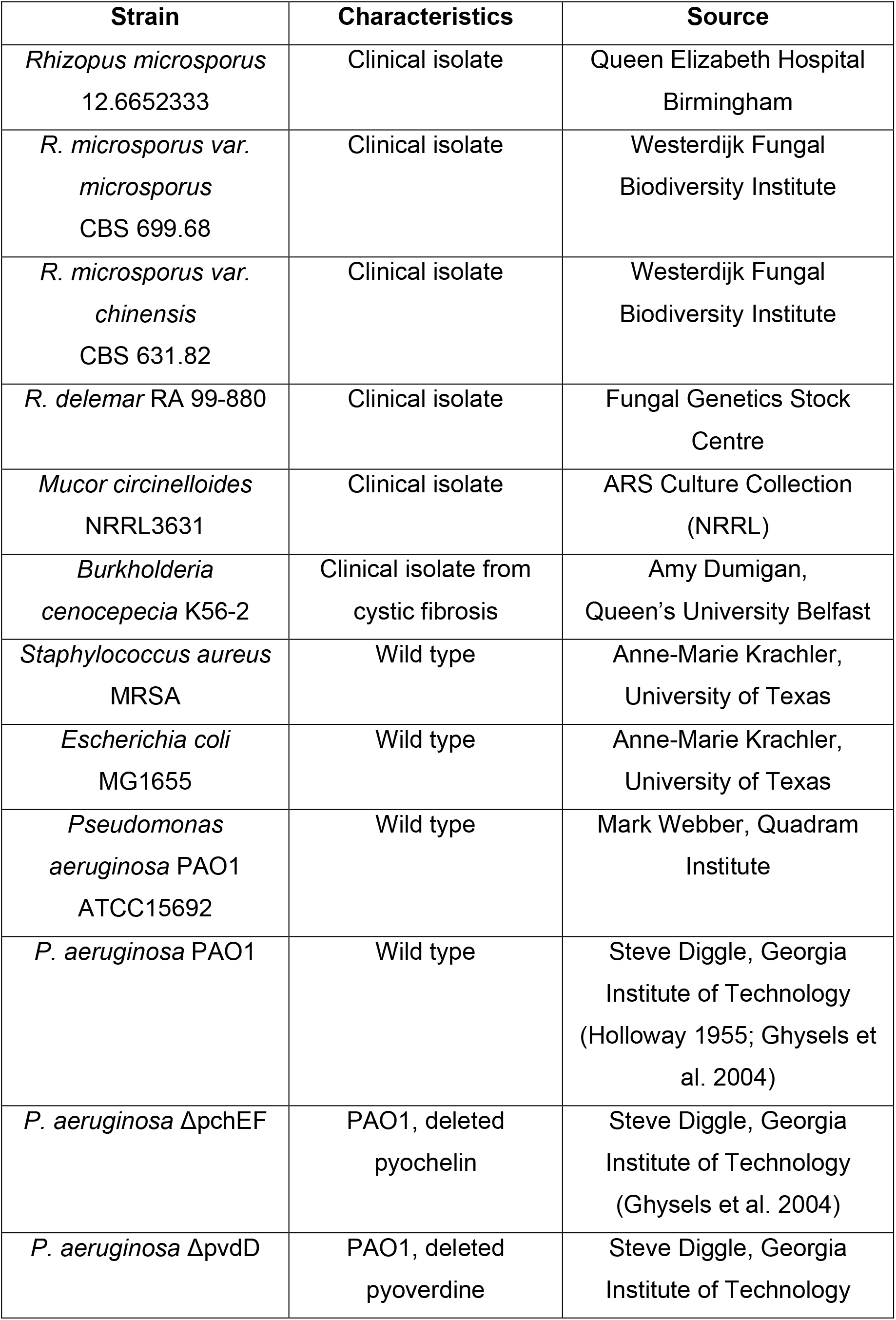
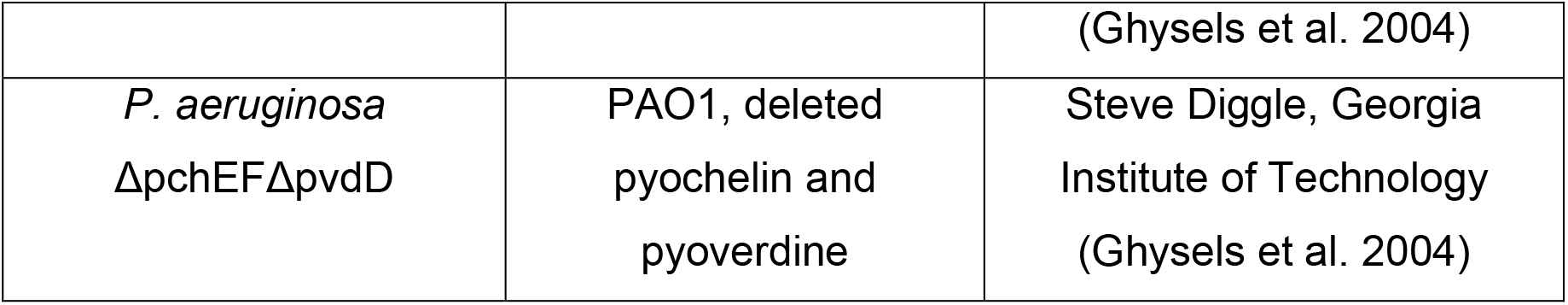
Strains used in this study.

### Live co-cultures

LB broth was inoculated with *P. aeruginosa*, *B. cenocepecia*, *E. coli*, or *S. aureus* and incubated for 24 hours (37°C, 200 rpm). Bacteria were washed three times with phosphate buffered solution (PBS). *R. microsporus* sporangiospores were harvested through flooding with PBS, washed once, and counted via haemocytometer. Spores (1 x 10^4^ spores/ml) were added to 50% SAB, 50% LB in a 96-well plate. Bacteria were added to each well at a multiplicity of infection (MOI) ratio of 1:1, 1:10, 1:50, and 1:100 and incubated for 24 h (static, 37°C). Wells were imaged using an inverted Zeiss AxioObserver microscope (20x magnification) and the number of germinated spores per field of view quantified. Germination was defined as the point in which the germ tube reached the same size as the spore diameter.

### Spore germination when exposed to bacterial supernatants

Bacterial cultures were prepared as previously detailed, and grown to at least stationary phase (OD_600_> 3.0). Cultures were centrifuged (3220 x g, 10 minutes) and the resulting supernatant filter sterilised. Sterile supernatants were stored at −80°C until required. Spores (1 x10^4^/ml) were added to a 96-well plate containing either 50% SAB and 50% LB broth, or 50% SAB and 50% supernatant. Fungal growth was determined by endpoint analysis using OD_600_ as a quantifier of growth (FLUOstar Omega plate reader).

To investigate the role of iron restriction, ferric chloride (100 mM) was diluted to 1, 10, 50, 100, 200, and 500 µM in supernatants. The iron was allowed to associate with any chelating molecules for 15 minutes before the addition of an equal volume of SAB. Wells containing the SAB/supernatant mixture without iron were included as controls.

### Live cell imaging

Live-cell imaging was performed for 12-18 hr at 37°C with humidity using a Zeiss AxioObserver microscope (20x magnification). Images were taken every 10 minutes to create a time-lapse movie, and the percentage of germinated spores in each field of view was determined.

### Exposure of pre-germinated spores to *P. aeruginosa* supernatant

Spores were harvested and added to 500 µl of SAB broth at a concentration of 1×10^6^ spores/ml in triplicate in a 24 well plate. Spores were incubated statically for 4-5 hours at 37°C until germlings emerged and then either 500 µl of *P. aeruginosa* supernatant or 500 µl LB was added. The plate was incubated for 18 h at 37°C, and the endpoint absorbance (OD_600_) of each well measured.

### Viability of spores exposed to *P. aeruginosa* supernatant

*R. microsporus* spores (1 x 10^6^ spores/ml) were exposed to 100% *P. aeruginosa* supernatant for 96 hr (statically, 37°C). Every 24 h, 100 spores were plated on SAB agar and incubated at 25°C for 24 h. Following incubation, the number of viable spores were counted and compared to 0 h control plates.

### Pyocyanin secretion

*P. aeruginosa* supernatants were prepared as described previously. Absorbance (690 nm) was measured using a FLUOstar Omega plate reader and compared to a pyocyanin standard curve.

### Organic extraction of *P. aeruginosa* siderophore mutant supernatants

Supernatants (WT and ΔpchEFΔpvdD) were mixed with Amberlite^®^ XAD-2^®^ resin for 1 h to permit binding of organic molecules to the resin. Samples were centrifuged (3,220 x g, 10 min) to pellet the resin and the resulting supernatant filter sterilised. LB was used as a procedural control.

### RNA extraction

*R. microsporus* spores (2.5 x 10^6^ spores/ml) were exposed to SAB/LB (50:50) (media only control) or 50% PAO1 supernatant. Flasks were incubated statically at 37°C for 7 h. Spores were centrifuged (1,811 x g, 3 min), snap frozen in liquid nitrogen, and stored at −80°C. When ready to extract RNA, 1ml of TRIzol (Invitrogen) was added to each sample and thawed on ice. These samples were homogenised as before. Chloroform (200 µl) was added to each sample, vortexed thoroughly, and centrifuged at 4°C, 9,400 x g for 15 min. The aqueous layer was collected and an equal volume of 100% ethanol was added. 700 µl of this was transferred to RNeasy columns, and the RNeasy Mini Plus Kit (Qiagen) protocol followed according to manufacturer guidelines. The RNA concentration and quality was measured using a spectrophotometer.

### Quantitative Reverse Transcriptase PCR

qRT-PCR was performed using an iTaq Universal SYBR Green One-Step Kit (Bio Rad) using 10ng/μl RNA with a total reaction volume of 20 μl. Protocol was followed according to manufacturer’s recommendations. *FTR1* was amplified using the forward primer (5’-GTGGTGTCTCCTTGGGTGTT-3’) and reverse primer (5’-CCACCACGGTAGATGAGGA-3’). This was normalised to 18s rRNA using the forward primer (5’-GGCGACGGTCCACTCGATTT3’) and reverse primer (5’-TCACTACCTCCCCGTGTCGG-3’).

### Quantification of overall siderophore production

Siderophore concentrations in bacterial supernatants were quantified by using the SideroTec Assay Kit (Emergen Bio) according to the manufacturer recommendations.

### Quantification of pyoverdine production in co-culture

*P. aeruginosa* and *R. microsporus* (1 x 10^6^ spores/ml) were co-cultured in 1 ml RPMI-1640 (Thermo-Fisher) at MOIs of 1:1, 1:10, 1:50, and 1:100 in a 24 well plate (24 h, 37°C). Plates were centrifuged (3,220 x g, 5 minutes) and 200 µl from each well was transferred to a 96 well plate in duplicate. Pyoverdine production was measured at 405 nm (Stintzi et al. 1996).

### Zebrafish infections

Adult wild type (AB) *Danio rerio* zebrafish were maintained at the University of Birmingham Aquatic Facility in recirculating tanks with 14 h light/10 h dark cycles at 28°C. Adult zebrafish naturally spawned overnight in groups of 11 fish (six female, five males). Embryos were transferred to E3 medium (5 mM NaCl, 0.17 mM KCl, 0.33 mM CaCl_2_, 0.33 mM MgSO_4_, pH 7) with 0.3 µg/ml methylene blue and 0.003% 1-phenyl-2-thiourea (PTU) for the first 24 hours post fertilisation (hpf) and maintained at 32°C.

Hindbrain injections were performed as previously described (Voelz et al. 2015). Sample sizes were calculated via power analysis using an alpha value of 0.05, power of 80%, mean effect size of 4.2%, and standard deviation of 8%, based on preliminary data and standards accepted by the zebrafish infection community. At 24 hpf larvae were manually dechorionated and anaesthetised (160 µg/ml Ethyl 3-aminobenzoate methanesulfonate salt [Tricaine]). *R. microsporus* spores were suspended in either polyvinylpyrrolidone (PVP, 10% in PBS + 0.05% phenol red) or PVP + 80 µg/ml pyoverdine at a concentration of 5 x 10^6^ spores/ml. Suspended spores (2 nl) were injected into the hindbrain via the otic vesicle to achieve a dose of 50 spores/larva. Control larvae were injected with either PVP only or PVP + 80 µg/ml pyoverdine. Any fish that did not survive the injection process were removed. Survival was recorded every 24 h until larvae were sacrificed at 5 dpf (96 hours post infection) through 10x overdose of Tricaine. Data were pooled from four separate experiments with a total of 87, 80, 80, and 81 fish for control, pyoverdine only, spores only, and spores + pyoverdine, respectively.

### Statistical analysis

Each experiment was performed with at least two technical and two biological replicates. Microsoft Excel 2016 and GraphPad Prism 6 were used to record and analyse data. Statistical tests used are indicated in figure legends. All analysis was performed on non-normalised raw data or arcsine transformed data where appropriate. A p-value of p < 0.05 was considered to indicate statistical significance. Statistical significance is indicated by * = p < 0.05, ** = p < 0.01, and *** = p < 0.001.

## Acknowledgements

We would like to acknowledge Steve Diggle, Anne-Marie Krachler, José Bengochea, Amy Dumigan, and Mark Webber for the generous gifts of bacterial strains; Francisco Fernandez-Trillo, Oliver Creese and Susan Moody for assistance with the organic extractions; Kevin Waldron and Daniel Stones for valuable consultation regarding experimental design with metals; Fabien Cottier for critical input while preparing the manuscript; Elizabeth Ballou for help with power calculations for animal studies; and the Host and Pathogen Interaction laboratory at the University of Birmingham for helpful discussion and valuable support.

C.K is funded by the Darwin Trust of Edinburgh. R.A.H and C.C are funded by the Medical Research Council (MR/L00903X/1).

---

**Video 1. *R. microsporus* germination in 50% SAB, 50% LB.**Images were taken every 10 min for 11 h using an inverted Zeiss AxioObserver microscope at 20x magnification. Scale bar represents 10 μM.

**Video 2. The presence of *P. aeruginosa* supernatant inhibits the germination of *R. microsporus* spores.** Spores were exposed to 50% *P. aeruginosa* supernatant and images were taken every 10 min for 11 h using an inverted Zeiss AxioObserver microscope at 20x magnification. Scale bar represents 10 μM.

## Figure supplement legends

**Figure 2 – figure supplement 1.**
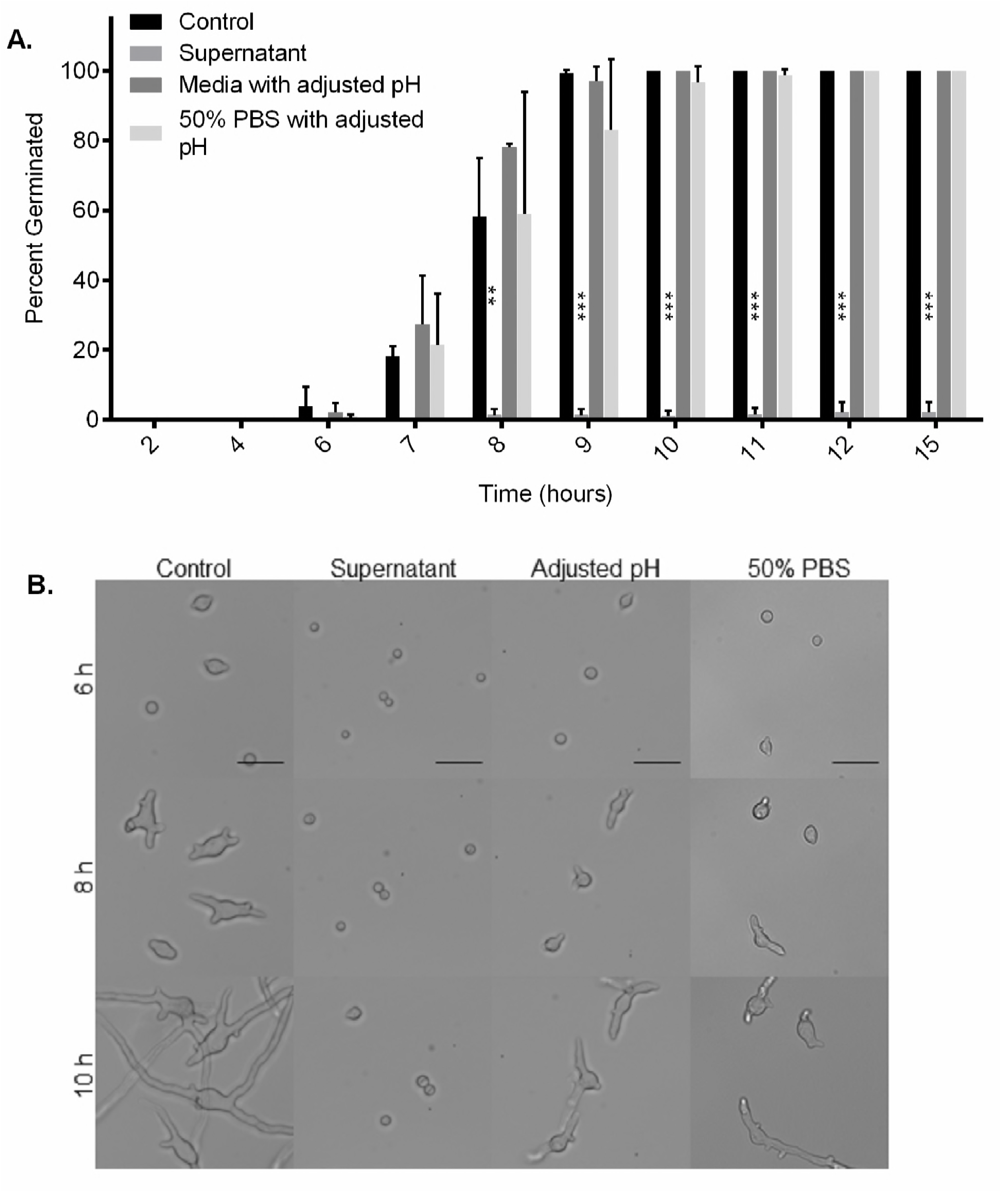
*Pseudomonas aeruginosa* inhibits the germination of *R. microsporus* independent of pH and nutrient availability. The pH of media containing 50% PAO1 supernatant is approximately 7.3. To determine whether the inhibitory effect is caused by this change, *R. microsporus* spores were exposed to 50/50 SAB/LB with pH adjusted to 7.3. A 50% PBS control was also used to determine effects of nutrient deprivation. These conditions were imaged over time and (A) the percent of *R. microsporus* spores germinated over time was quantified and (B) representative images were obtained (n=2, Two-way ANOVA performed on arcsine transformed data). Scale bars = 50 µm. Error bars depict SEM. Source files for all microscopy movies used in the quantitative analysis are available in Figure 2 – figure supplement 1-Source data 1.

**Figure 3 – figure supplement 1.**
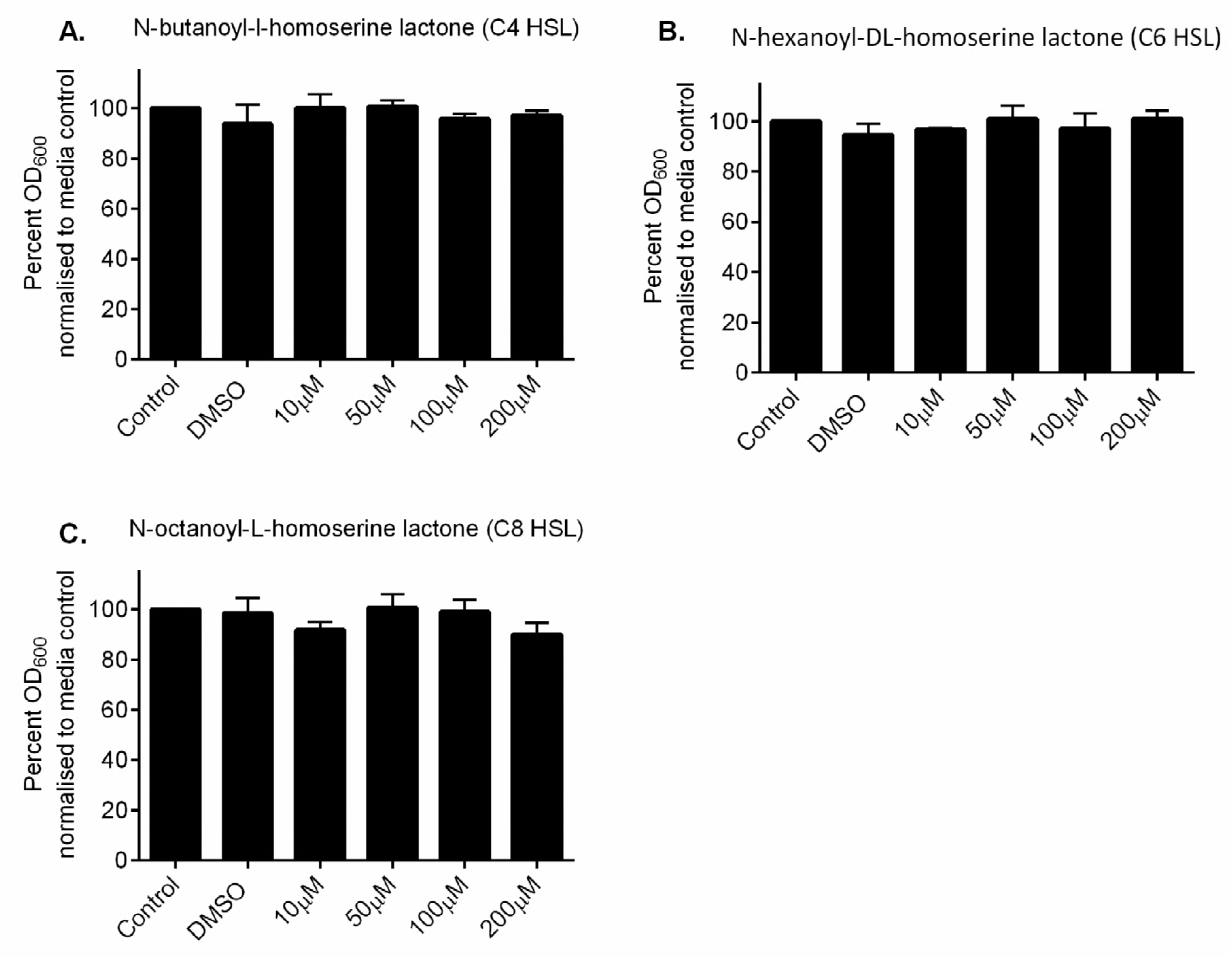
Inhibition of *R. microsporus* germination is not mediated by quorum sensing molecules or toxin exposure. *R. microsporus* spores were incubated in SAB media for 24 h at 37°C with increasing concentrations of (A) N-butanoyl-l-homoserine lactone (C4 HSL), (B) N-hexanoyl-DL-homoserine lactone (C6 HSL), (C) N-octanoyl-L-homoserine lactone (C8 HSL). Fungal growth was quantified by absorbance (OD_600_) and normalised to media control (n=3). All data was analysed by a Kruskal-Wallis test with Dunn’s multiple comparisons test. Error bars depict SEM.

**Figure 4 – figure supplement 1.**
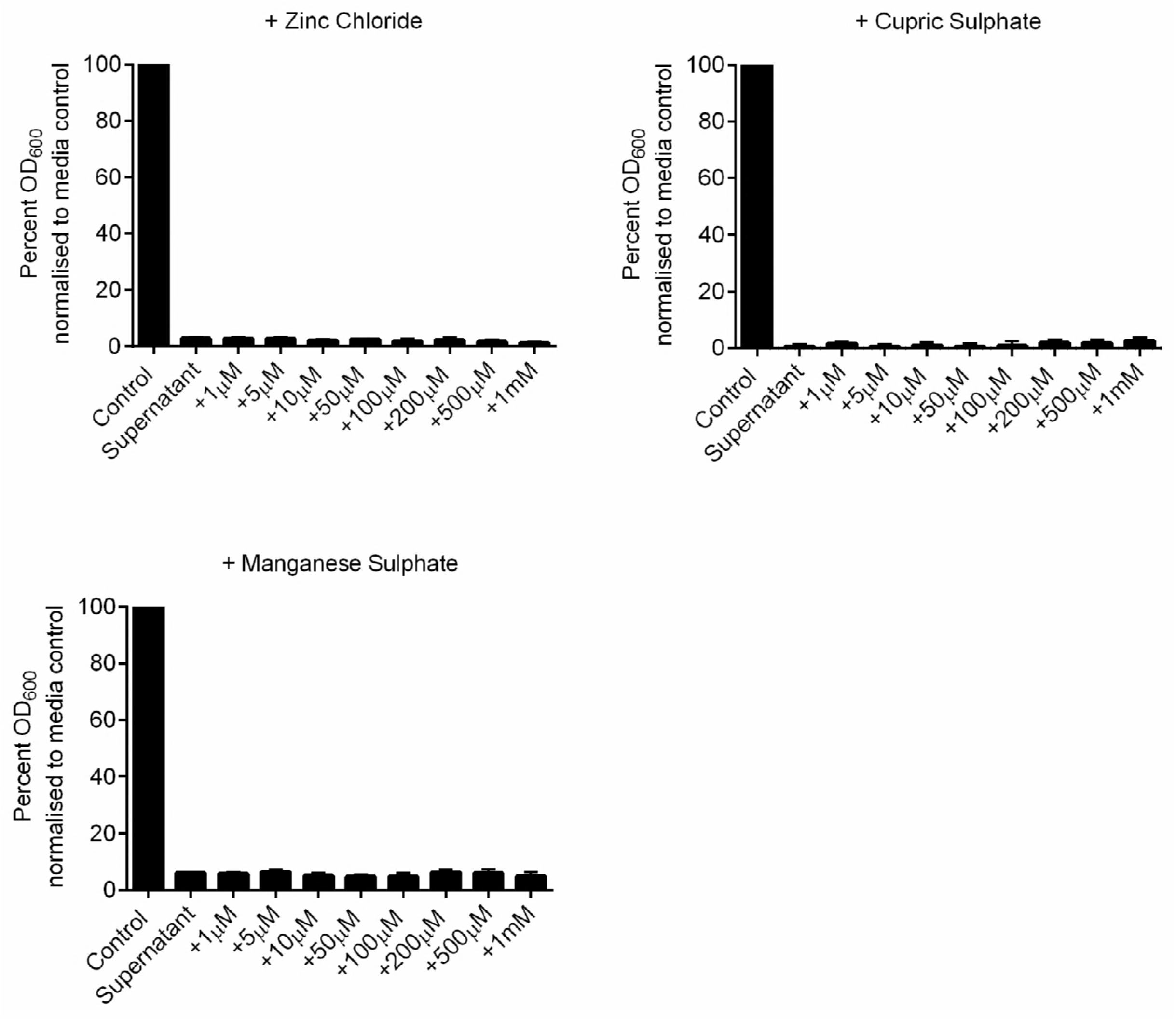
Inhibition of *R. microsporus* growth by *P. aeruginosa* supernatant is not due to zinc, copper, or manganese restriction. *R. microsporus* spores were exposed to 50% *P. aeruginosa* supernatant and spiked with increasing concentrations of zinc chloride, cupric sulphate, and manganese sulphate for 24 h statically at 37°C. Fungal growth was measured through absorbance (OD_600_) and normalised to media control (n=3, Kruskal-Wallis with Dunn’s multiple comparisons test).

## Data source legends

**Figure 1-Source Data file 1. Source data files for *R. microsporus* + *P. aeruginosa* co-culture.** This zip folder contains DIC images taken after 24-hour co-culture, using a Nikon TE2000-U inverted microscope with 20x objective. Each folder is labelled by the experiment number and date performed. These are further divided into sub-folders labelled by experimental condition. “MOI 1” indicates a multiplicity of infection ratio 1:1, “MOI 10” indicates a multiplicity of infection ratio 1:10, etc. SABLB Control indicates that the spores were grown in fresh 50% SAB, 50% LB.

The files were exported as 16-bit .tif files (1280 x 1024 pixel resolution, 0.29µm/pixel). Please note, these images must be opened using a program, such as ImageJ. Scale bars depict 20 µm.

**Figure 2-source data 1. Source files for live cell imaging of spores germinating over time when exposed to *P. aeruginosa* supernatant.** This zip folder contains DIC images taken every 10 minutes over 18 hours to create a movie in real-time. Each folder is labelled by the experiment number and date performed. These are further divided into subfolders labelled by experimental condition. “Media control” indicates that the spores were grown in fresh 50% SAB, 50% LB. “50% PAO1 supernatant” indicates that the spores were grown in 50% *P. aeruginosa* supernatant, 50% SAB.

Images were taken using a Zeiss AxioObserver inverted microscope with 20x long working distance objective in an environmental control chamber (37°C, no CO_2_, humidity).

The files were exported as 16-bit .czi files (2048 x 2048 pixel resolution, 0.315 µm/pixel). As these files must be opened using ImageJ Fiji, these image stacks were converted to .avi at 5 frames per second, which can be viewed using common media viewers. These are organised into folders labelled “CZI files” or “AVI files,” respectively.

Each movie begins at a different timepoint, as it takes a varying amount of time to set up the microscope each time. “Expt 1 09.05.16” begins at 30 minutes, “Expt 2 17.05.16” begins at 25 minutes, “Expt 3 19.05.16” begins at 20 minutes, and “Expt 4 12.07.16” begins at 25 minutes. Scale bars depict 50 µm.

**Figure 4-source data 1. Source data files for *R. microsporus*and *P. aeruginosa* co-culture and iron.** This zip folder contains DIC images taken after 24-hour co-culture using a Nikon TE2000-U inverted microscope with 20x objective.

Each folder is labelled by the experiment number and date performed. These are further divided into sub-folders labelled by experimental condition. “MOI 1” indicates a multiplicity of infection ratio 1:1, “MOI 10” indicates a multiplicity of infection ratio 1:10, etc. SABLB Control indicates that the spores were grown in fresh 50% SAB, 50% LB.

The files were exported as 16-bit .tif files (1280 x 1024 pixel resolution, 0.29µm/pixel). Please note, these images must be opened using a program, such as ImageJ. Scale bars depict 50 µm.

**Figure 2 –figure supplement 1-Source data 1. Source files for live cell imaging of spores germinating over time when exposed to 50% *P. aeruginosa* supernatant, SAB/LB with a pH adjusted to 7.3.** This zip folder contains DIC images taken every 10 minutes over 18 hours to create a movie in real-time. Each folder is labelled by the experiment number and date performed. These are further divided into subfolders labelled by experimental condition. “50% PAO1 supernatant” indicates that the spores were grown in 50% *P. aeruginosa* supernatant, 50% SAB. “50% PBS” indicates that the spores were grown in 50% PBS, 50% SAB. “pH adjusted SAB.LB” indicates that the spores were exposed to 50% SAB, 50% LB with a pH of 7.3. “SAB.LB control” indicates that the spores were grown in fresh 50% SAB, 50% LB.

Images were taken using a Zeiss AxioObserver inverted microscope with 20x long working distance objective in an environmental control chamber (37°C, no CO_2_, humidity).

The files were exported as 16-bit .czi files (2048 x 2048 pixel resolution, 0.315 µm/pixel). As these files must be opened using ImageJ Fiji, these image stacks were converted to .avi at 5 frames per second, which can be viewed using common media viewers. These are organised into folders labelled “CZI files” or “AVI files,” respectively.

Each movie begins at a different timepoint, as it takes a varying amount of time to set up the microscope each time. “Expt 1 08.09.16” begins at 30 minutes, and “Expt 2 16.08.16” begins at 50 minutes.

